# Alarming antibody evasion properties of rising SARS-CoV-2 BQ and XBB subvariants

**DOI:** 10.1101/2022.11.23.517532

**Authors:** Qian Wang, Sho Iketani, Zhiteng Li, Liyuan Liu, Yicheng Guo, Yiming Huang, Anthony D. Bowen, Michael Liu, Maple Wang, Jian Yu, Riccardo Valdez, Adam S. Lauring, Zizhang Sheng, Harris H. Wang, Aubree Gordon, Lihong Liu, David D. Ho

## Abstract

The SARS-CoV-2 Omicron variant continues to evolve, with new BQ and XBB subvariants now rapidly expanding in Europe/US and Asia, respectively. As these new subvariants have additional spike mutations, they may possess altered antibody evasion properties. Here, we report that neutralization of BQ.1, BQ.1.1, XBB, and XBB.1 by sera from vaccinees and infected persons was markedly impaired, including sera from individuals who were boosted with a WA1/BA.5 bivalent mRNA vaccine. Compared to the ancestral strain D614G, serum neutralizing titers against BQ and XBB subvariants were lower by 13-81-fold and 66-155-fold, respectively, far beyond what had been observed to date. A panel of monoclonal antibodies capable of neutralizing the original Omicron variant, including those with Emergency Use Authorization, were largely inactive against these new subvariants. The spike mutations that conferred antibody resistance were individually studied and structurally explained. Finally, the ACE2-binding affinities of the spike proteins of these novel subvariants were found to be similar to those of their predecessors. Taken together, our findings indicate that BQ and XBB subvariants present serious threats to the efficacy of current COVID-19 vaccines, render inactive all authorized monoclonal antibodies, and may have gained dominance in the population because of their advantage in evading antibodies.

## INTRODUCTION

The coronavirus disease 2019 (COVID-19) pandemic, caused by severe acute respiratory syndrome coronavirus 2 (SARS-CoV-2), continues to rage due to emergence of the Omicron variant and its descendant subvariants (Iketani et al., 2022; Liu et al., 2022b; Wang et al., 2022d; Wang et al., 2022e; Wang et al., 2022f). While the BA.5 subvariant is globally dominant at this time (**Figure 1A**), a diverse array of Omicron sublineages have arisen and are competing in the so-called “variant soup” (Callaway, 2022). It has become apparent that four new subvariants are rapidly gaining ground on BA.5, raising the specter of yet another wave of infections in the coming months. BQ.1 and BQ.1.1 were first identified in Nigeria in early July and then expanded dramatically in Europe and North America, now accounting for 46%, 32%, and 44% of cases in France, the United Kingdom, and the United States, respectively (**Figure 1A**). XBB and XBB.1 were first identified in India in mid-August and quickly became predominant in India, Singapore, and other regions in Asia (**Figure 1A**). BQ.1 and BQ.1.1 evolved from BA.5, whereas XBB and XBB.1 resulted from a recombination between two BA.2 lineages, BJ.1 and BA.2.75 (**Figure 1B**). The spike protein of BQ.1 harbors the K444T and N460K mutations in addition to those found in BA.5, with BQ.1.1 having an additional R346T mutation (**Figures 1C and S1**). Strikingly, the spike of XBB has 14 mutations in addition to those found in BA.2, including 5 in the N-terminal domain (NTD) and 9 in the receptor-binding domain (RBD), whereas XBB.1 has an additional G252V mutation (**Figures 1C and S1**). The rapid rise of these subvariants and their extensive array of spike mutations are reminiscent of the appearance of the first Omicron variant last year, thus raising concerns that they may further compromise the efficacy of current COVID-19 vaccines and monoclonal antibody therapeutics. We now report findings that indicate that such concerns are, sadly, justified, especially so for the XBB and XBB.1 subvariants.

**Figure 1.**
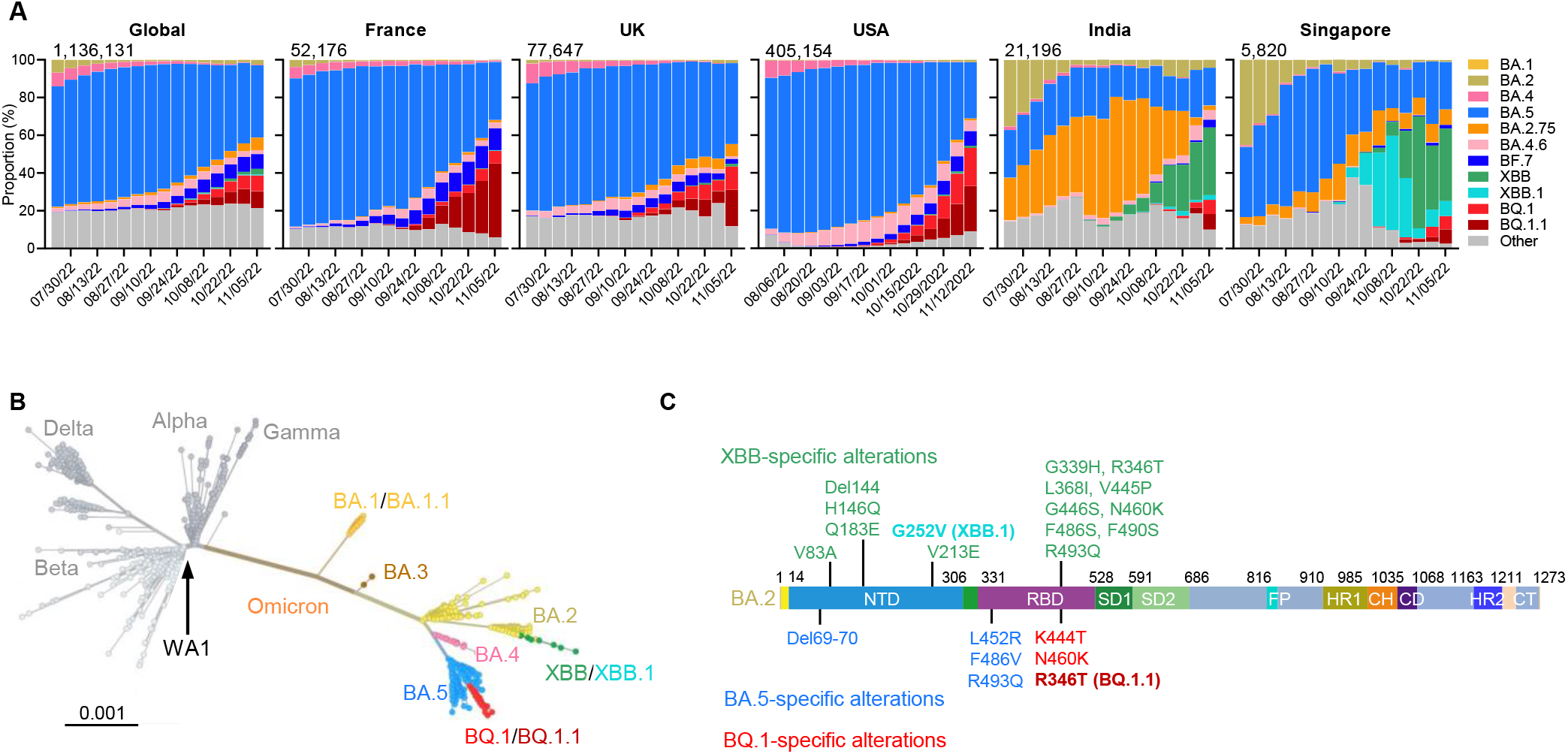
The rise of SARS-CoV-2 Omicron BQ.1, BQ.1.1, XBB, and XBB.1 subvariants. (A) Frequencies of Omicron subvariants from the CDC (USA) or GISAID (all other countries). The values in the upper left corner of each box denotes the cumulative number of sequences for all circulating viruses in the denoted time period. (B) Unrooted phylogenetic tree of Omicron subvariants along with other main SARS-CoV-2 variants. The scale bar indicates the genetic distance. (C) Key spike mutations found in XBB and XBB.1 in the background of BA.2 and in BQ.1 and BQ.1.1 in the background of BA.4/5. Del, deletion. The positions of these mutations on the spike trimer are shown in **Figure S1**.

## RESULTS

### Neutralization by polyclonal sera

To understand if BQ.1, BQ.1.1, XBB, and XBB.1 have stronger resistance to serum antibodies, we first set out to evaluate the neutralization of these four new subvariants by sera from five different clinical cohorts. These results are summarized in **Figure 2**. The five clinical cohorts included individuals who received three or four doses of one of the original COVID-19 mRNA vaccines (termed “3 shots WT” or “4 shots WT”, respectively), those who received one of the recently authorized bivalent (WT and BA.5) COVID-19 mRNA vaccines as a 4^th^ shot after three doses of one of the original COVID-19 mRNA vaccines (termed “3 shots WT + bivalent”), and patients who had BA.2 and BA.4 or BA.5 breakthrough infection after vaccination (termed “BA.2 breakthrough” and “BA.4/5 breakthrough”, respectively). Their relevant clinical information is summarized in **Table S1**. Consistent with previous findings (Wang et al., 2022d), BA.2 and BA.4/5 showed stronger evasion to serum neutralization relative to the ancestral strain D614G across all five cohorts (**Figure 2A**). The geometric mean 50% inhibitory dose (ID_50_) titers against BA.2 and BA.4/5 decreased 2.9-to 7.8-fold and 3.7-to 14-fold, respectively, compared to that against D614G. Alarmingly, in the “3 shots WT” cohort, neutralization titers were far lower against BQ.1, BQ.1.1, XBB, and XBB.1, with reductions of >37-fold to >71-fold compared to D614G. Moreover, while all sera had detectable titers against BA.2 and BA.4/5, a majority of samples did not neutralize the new subvariants at the lowest dilution (1:100) of serum tested. A similar trend was also noted in the other four cohorts, with the lowest titers observed against XBB.1, followed by XBB, BQ.1.1, and BQ.1. The geometric mean neutralization titers of sera from the “BA.4/5 breakthrough” and “BA.2 breakthrough” cohorts were noticeably higher, indicating that SARS-CoV-2 breakthrough infection induced better antibody responses than vaccination among these samples.

**Figure 2.**
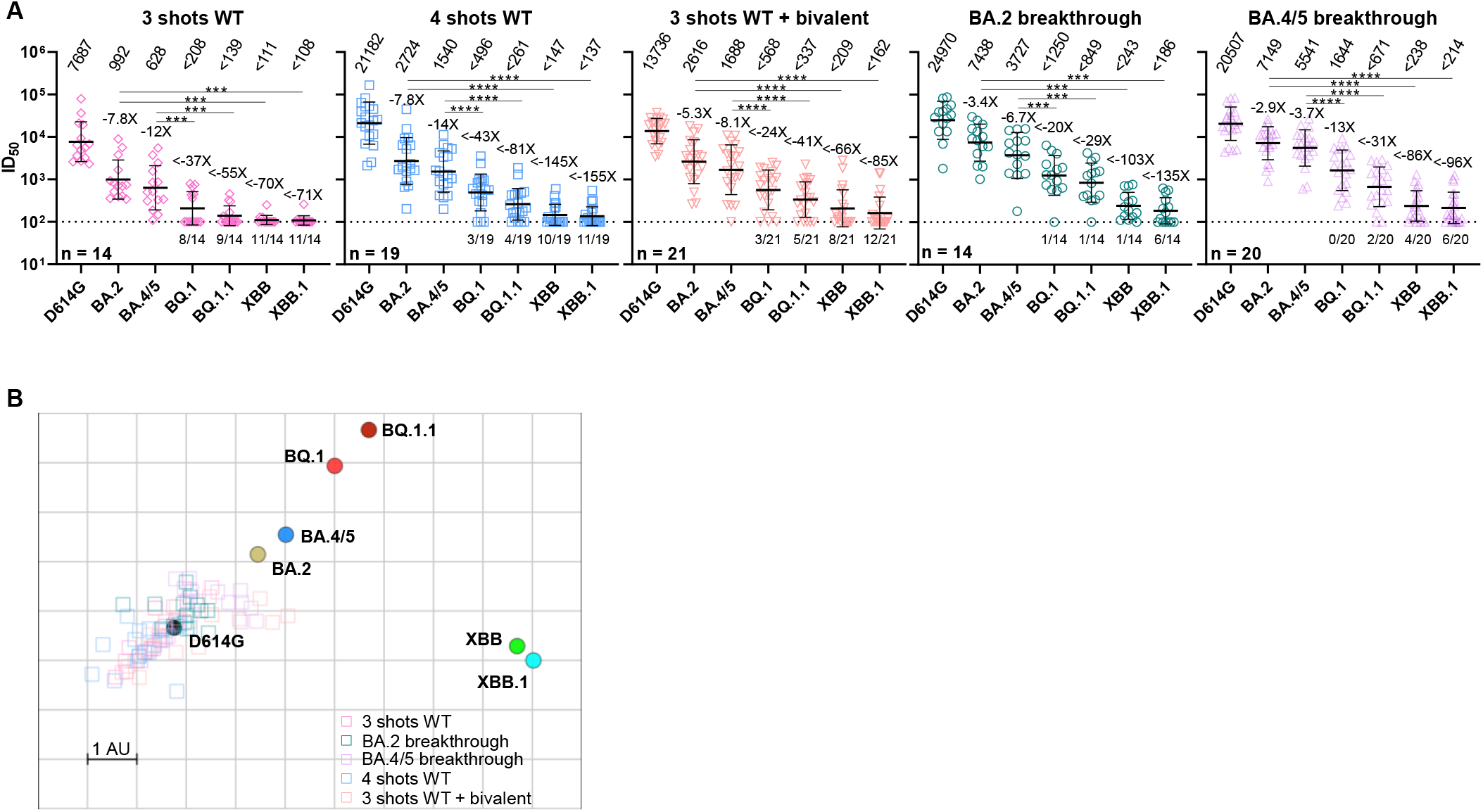
Serum neutralization of Omicron subvariants BQ.1, BQ.1.1, XBB, and XBB.1. (A) Neutralization of pseudotyped D614G and Omicron subvariants by sera from five different clinical cohorts, with their clinical information summarized in Table S1. The limit of detection is 100 (dotted line). Values above the symbols denote the geometric mean ID_50_ values, and values beneath the symbols denote the numbers of samples that lost neutralization activity. Values on the lower left show the sample size (*n*) for each group. The fold reduction in geometric mean ID_50_ value for each variant compared to D614G is also shown above the symbols. Comparisons were made by two-tailed Wilcoxon matched-pairs signed-rank tests. \*\*\**p* < 0.001; \*\*\*\**p* < 0.0001. (B) Antigenic map based on the serum neutralization data from (A). Virus positions are represented by closed circles while serum positions are shown as open squares. Sera are colored by group. Both axes represent antigenic distance with one antigenic distance unit (AU) in any direction corresponding to a two-fold change in neutralization ID_50_ titer.

We then utilized the serum neutralization results to construct an antigenic map to depict the antigenic distances among D614G and the Omicron subvariants (Smith et al., 2004) (**Figure 2B**). The resulting map shows that BQ.1.1 has drifted away from BA.4/5 antigenically as much as the latter has from the ancestral D614G. With each antigenic unit equaling a 2-fold difference in virus neutralization, BQ.1.1 is approximately 6-fold more resistant to serum neutralization than its predecessor BA.5. On the other hand, it is clear that XBB.1 is the most antigenically distinct of the Omicron subvariants. The large number of antigenic units that separates XBB.1 and BA.2 suggests that this new subvariant is ~63-fold more resistant to serum neutralization than its predecessor, or ~49-fold more resistant than BA.4/5. The impact of this antigenic shift on vaccine efficacy is particularly concerning.

### Neutralization by monoclonal antibodies

To understand the types of serum antibodies that lost neutralizing activity against BQ.1, BQ.1.1, XBB, and XBB.1, we constructed pseudoviruses for each subvariant, as well as for each individual mutation found in the subvariants, and then evaluated their susceptibility to neutralization by a panel of 23 monoclonal antibodies (mAbs) targeting various epitopes on the spike (**Figure 3A**). These mAbs were chosen because they had appreciable activity against the initial Omicron variant. Among these antibodies, 20 were directed to the class 1 to class 4 epitope clusters on the RBD (Barnes et al., 2020): S2K146 (Park et al., 2022), Omi-3 (Nutalai et al., 2022), Omi-18 (Nutalai et al., 2022), BD-515 (Cao et al., 2021), XGv051 (Wang et al., 2022b), XGv347 (Wang et al., 2022a), ZCB11 (Zhou et al., 2022), COV2-2196 (tixagevimab) (Zost et al., 2020), LY-CoV1404 (bebtelovimab, authorized to treat COVID-19) (Westendorf et al., 2022), XGv289 (Wang et al., 2022a), XGv264 (Wang et al., 2022b), S309 (sotrovimab) (Pinto et al., 2020), P2G3 (Fenwick et al., 2022), SP1-77 (Luo et al., 2022), BD55-5840 (Cao et al., 2022), XGv282 (Wang et al., 2022a), BD-804 (Du et al., 2021), 35B5 (Wang et al., 2022g), COV2-2130 (cilgavimab) (Zost et al., 2020), and 10-40 (Liu et al., 2022a). The other three were non-RBD mAbs, with C1520 (Wang et al., 2022h) targeting the NTD, C1717 (Wang et al., 2022h) targeting NTD-SD2, and S3H3 (Hong et al., 2022) targeting SD1. We also included the clinical mAb combination of COV2-2196 and COV2-2130, marketed as Evusheld for the prevention of SARS-CoV-2 infection. Their neutralization IC_50_ values are presented in the **Figure S2** and their fold changes in IC_50_ compared to BA.4/5 or BA.2 are shown in **Figure 3B**. BQ.1 and BQ.1.1 were greatly or completely resistant to all RBD class 1 and class 3 mAbs tested as well as to one RBD class 2 mAb (XGv051), a class 4 mAb (10-40), and an NTD-SD2 mAb (C1717). The loss of neutralizing activity of NTD-SD2 and RBD class 1 mAbs were due to the N460K mutation, while the impairment in the potency of RBD class 3 mAbs resulted from both the R346T and K444T mutations. As BQ.1.1 has one more mutation (R346T) than BQ.1, it exhibited stronger antibody evasion to the class 3 RBD mAbs than BQ.1. Importantly, clinically authorized LY-CoV1404 (bebtelovimab) and Evusheld were inactive against BQ.1 or BQ.1.1.

**Figure 3.**
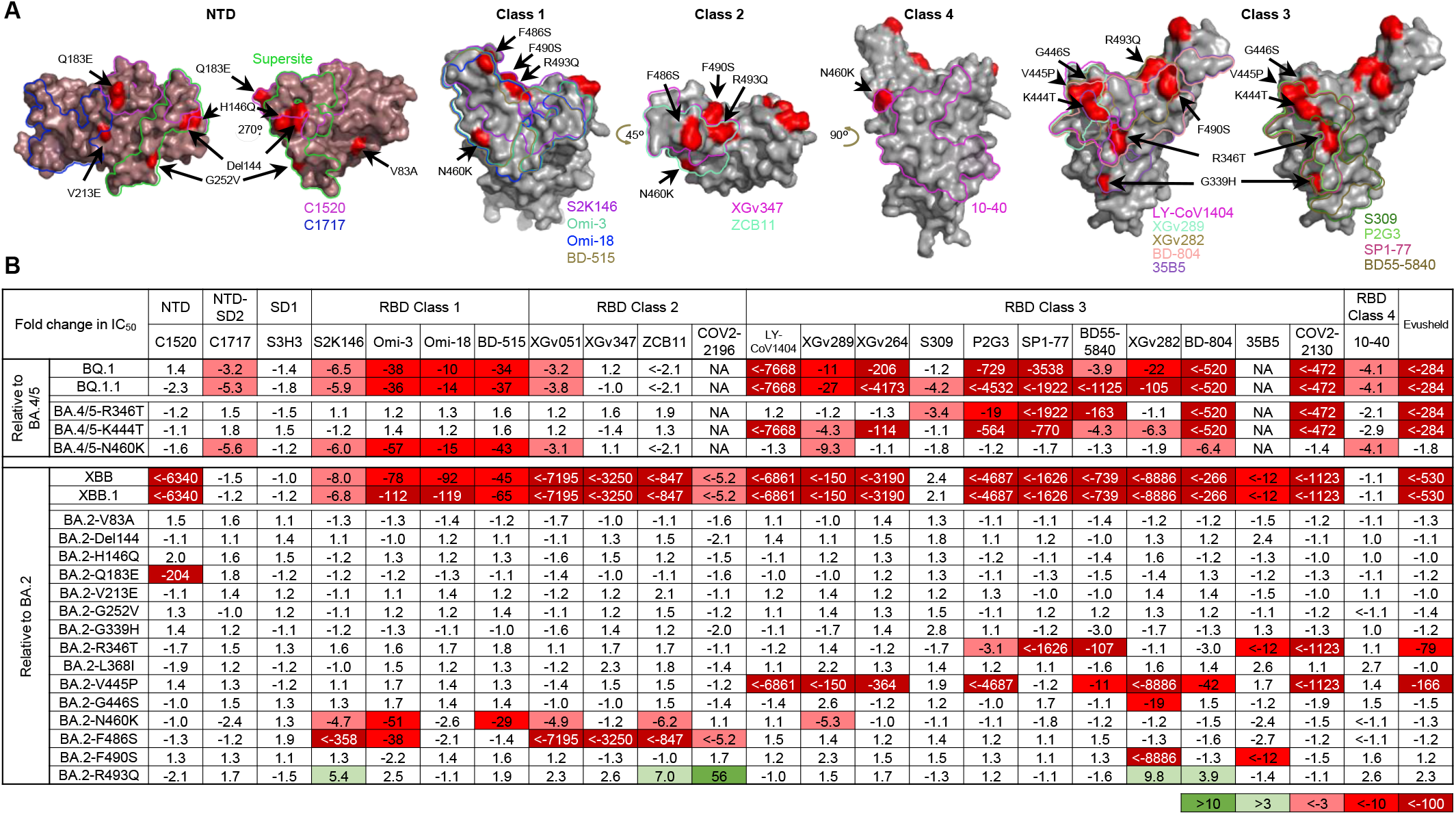
Resistance of Omicron subvariants to monoclonal antibody neutralization. (A) Footprints of NTD- and RBD-directed antibodies tested are outlined, and mutations within BQ.1, BQ.1.1, XBB, and XBB.1 are highlighted in red. (B) The fold changes in neutralization IC_50_ values of BQ.1, BQ.1.1, XBB, XBB.1, and the individual mutants compared with BA.4/5 or BA.2, with resistance colored red and sensitization colored green. The raw IC_50_ values are shown in **Figure S2**.

Against XBB and XBB.1, 19 of 23 mAbs lost neutralizing activity greatly or completely. Only C1717, S3H3, S309 (sotrovimab), and 10-40 showed relatively little fold change in neutralizing activity against these two subvariants relative to BA.2, although we note that these mAbs, with the exception of S3H3, had already lost significant activity against BA.2 relative to D614G (**Figure S2**). The Q183E mutation contributed to the activity loss of C1520; N460K and F486S accounted for the resistance to the RBD class 1 and class 2 mAbs; and R346T, V455P, G446S, and F490S contributed to the resistance to the RBD class 3 mAbs. Again, the clinically authorized LY-CoV1404 (bebtelovimab) and Evusheld could not neutralize XBB or XBB.1.

Several aforementioned point mutants (R346T, N460K, and F486S) had been observed in prior SARS-CoV-2 variants, and their impact on mAb binding have been reported (Wang et al., 2022d; Wang et al., 2022e; Wang et al., 2022f). We therefore conducted structural modeling to understand the impact of the newly identified point mutants (Q183E, K444T, V445P, and F490S) on the binding of select mAbs (**Figure 4**). The Q183E mutation in XBB and XBB.1 disrupted the hydrogen bond that residue A32 of mAb C1520 has with the spike and caused a steric clash with residue W91, likely abrogating the binding of this mAb (**Figure 4A**). K444T, found in BQ.1 and BQ.1.1, impaired the neutralization activities of most of the class 3 mAbs tested (**Figure 3B**), probably because mutating lysine to threonine made the side chain shorter and uncharged, which in turn would impair the interactions of this residue with mAbs directed to this site, as can be seen with SP1-77 and LY-CoV1404 (**Figures 4B and 4C**). Similarly, the V445P substitution in XBB and XBB.1 could exert an equivalent effect as K444T, by causing steric hindrance and/or disrupting a hydrogen bond with mAbs, resulting in the loss of antibody neutralization (**Figures 4D and 4E**). Finally, F490S impaired the neutralizing activities of XGv282, which can be accounted for by the abolition of a cation-π interaction (**Figure 4F**).

**Figure 4.**
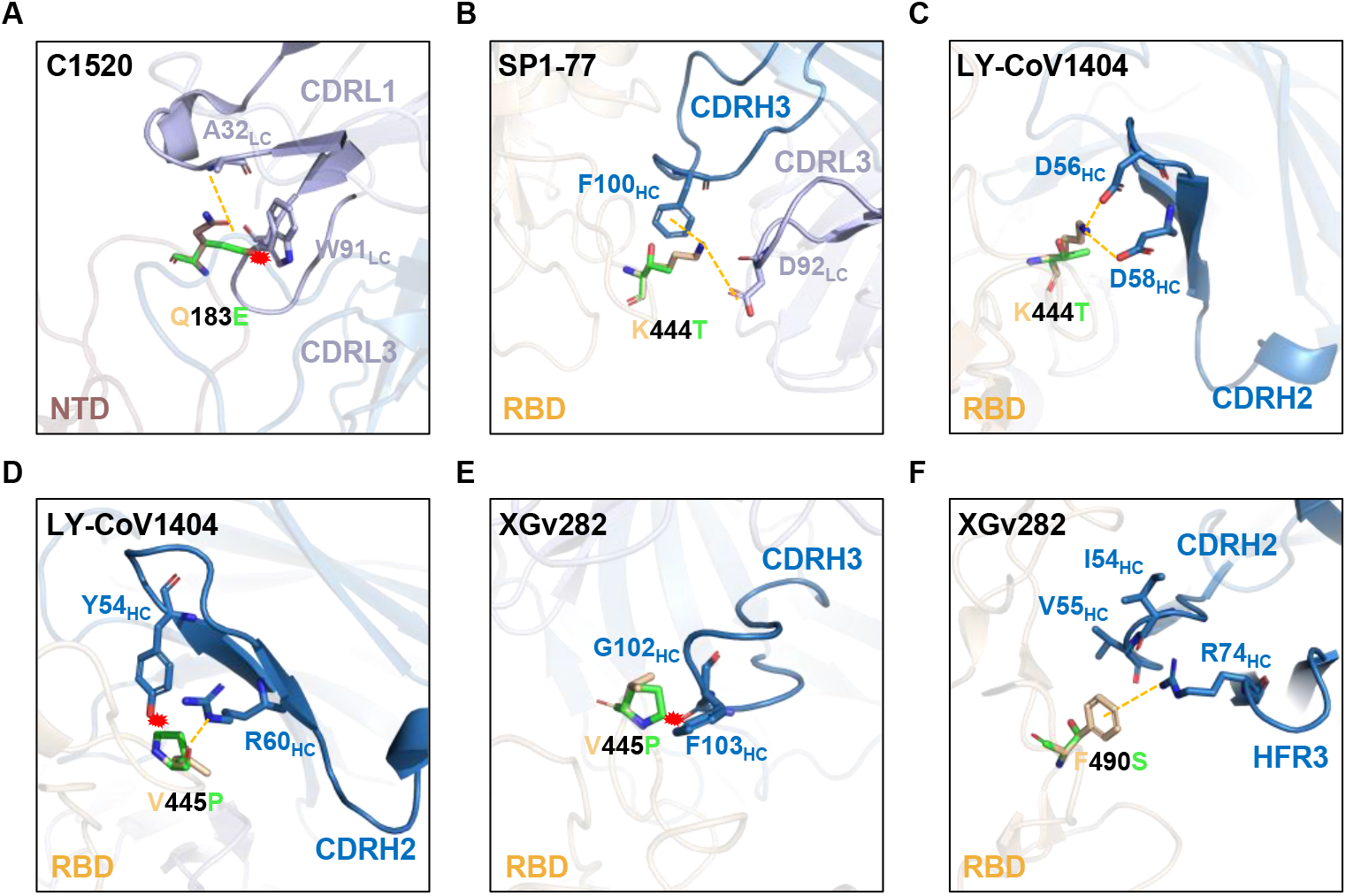
Structural analysis of mutational effects on binding of mAbs. Modeling of how (A) Q183E affects mAb C1520 neutralization, and how (B, C) K444T, (D, E) V445P, and (F) F490S affect RBD class 3 mAbs. Interactions are shown as yellow dotted lines and clashes are indicated as red asterisks.

### Receptor affinity

Angiotensin converting enzyme 2 (ACE2) is the receptor responsible for the entry of SARS-CoV-2 into target cells, and the binding affinity for this receptor may influence the transmissibility of the virus. We generated the spike trimer proteins of BA.2, BA.4/5, BQ.1, BQ.1.1, XBB, and XBB.1, and then tested their binding affinities to human ACE2 (hACE2) using surface plasmon resonance (SPR) (**Figure 5**). Our results showed that the viral receptor affinities of BQ.1 and BQ.1.1 spikes were comparable to that of BA.4/5 spike, with K_D_ ranging from 0.56 nM to 0.62 nM. The binding affinities for hACE2 of XBB and XBB.1 spikes exhibited a modest drop relative to that of BA.2 spike (K_D_ of 2.00 nM and 2.06 nM versus 0.95 nM). These findings suggested that the combination of mutations found in BQ.1 and BQ.1.1 did not alter the spike binding affinity to hACE2. The modest loss in hACE2 affinity for XBB and XBB.1 spikes may be due to F486S and R493Q mutations, which reside at the top of the RBD, where similar mutations, F486V and R493Q, were previously observed in BA.4/5 to impair and improve hACE2 binding, respectively (Wang et al., 2022d). In XBB and XBB.1, the serine rather than a valine may lower hACE2 binding, as has been observed in a deep mutational scanning study (Starr et al., 2022). Overall, these SPR measurements provide no evidence that the rise of these new subvariants is due to a higher affinity for hACE2.

**Figure 5.**
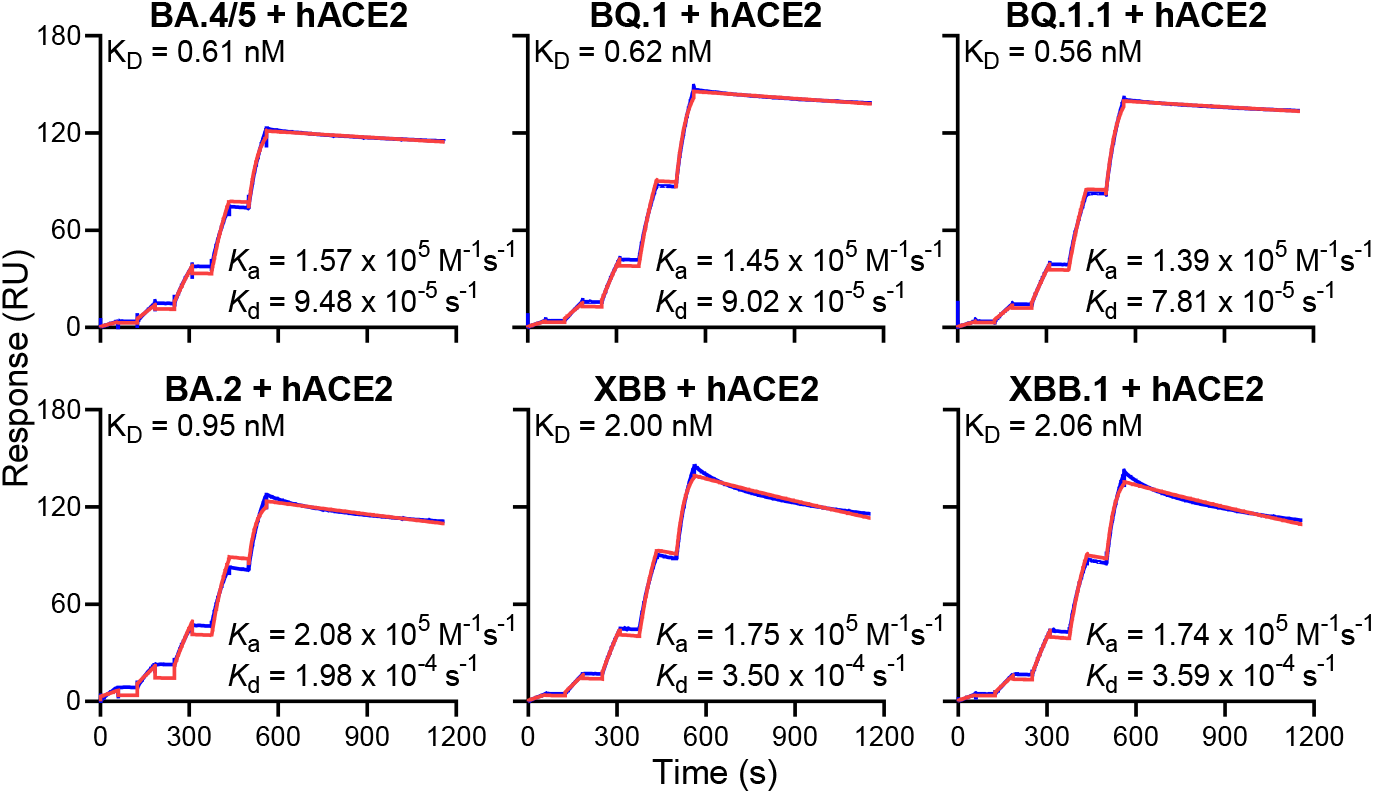
Receptor binding affinities of Omicron subvariant spikes. Each spike was produced and purified as prefusion-stabilized trimers, and their binding to human ACE2 was measured by SPR.

## DISCUSSION

In summary, we have examined in detail the antibody resistance profile and viral receptor binding affinity of SARS-CoV-2 Omicron BQ.1, BQ.1.1, XBB, and XBB.1 subvariants, which are rapidly expanding globally and already predominant regionally (**Figure 1A**). Our data demonstrate that these new subvariants were barely susceptible to neutralization by sera from vaccinated individuals with or without prior infection, including persons recently boosted with the new bivalent (WA1-BA.5) mRNA vaccines (**Figure 2**). The extent of the antigenic drift or shift measured herein is comparable to the antigenic leap made by the initial Omicron variant from its predecessors one year ago. In fact, combining these results with our prior findings on the serum neutralization of select sarbecoviruses (Wang et al., 2022c), there are indications that XBB and XBB.1 are now antigenically more distant than SARS-CoV or some sarbecoviruses in animals (**Figure S3**). Therefore, it is alarming that these newly emerged subvariants could further compromise the efficacy of current COVID-19 vaccines and result in a surge of breakthrough infections, as well as re-infections.

We also showed that these new subvariants were completely or partially resistant to neutralization by most monoclonal antibodies tested, including those with Emergency Use Authorization (**Figures 3B and S2**). These findings helped to define the causes behind the loss of serum neutralizing activity. BQ.1 and BQ.1.1 are largely pan-resistant to antibodies targeting the RBD class 1 and class 3 epitopes, whereas XBB and XBB.1 are pan-resistant to antibodies targeting the RBD class 1, 2, and 3 epitopes. These BQ and XBB sublineages have evolved additional mutations that are seemingly “filling up the holes” that allow a few mAbs to get through and neutralize their Omicron predecessors. Interestingly, both sublineages have converged on identical (R346T and N460K) or similar solutions (K444T versus V445P and G446S) to enhance antibody evasion. Furthermore, we have provided structural explanations for antibody resistance of various point mutants, including three that were previously undescribed (Q183E, K444T, and V445P) (**Figure 4**).

Perhaps the most important outcome of these mAb studies is the clinical implication for the use of mAbs to treat or prevent COVID-19. Previous SARS-CoV-2 variants have already successively knocked out the use of clinically authorized therapeutic antibodies (bamlanivimab, etesevimab, imdevimab, casirivimab, tixagevimab, cilgavimab, and sotrovimab), with bebtelovimab remaining as the only active monoclonal antibody against circulating SARS-CoV-2 strains (Iketani et al., 2022; Liu et al., 2022b; Wang et al., 2021; Wang et al., 2022d; Wang et al., 2022e; Wang et al., 2022f). Unfortunately, both BQ and XBB sublineages are now completely resistant to bebtelovimab, leaving us with no authorized antibody for treatment use. In addition, the combination of mAbs known as Evusheld that is authorized for the prevention of COVID-19 is also completely inactive against the new subvariants. This poses a serious problem for millions of immunocompromised individuals who do not respond robustly to COVID-19 vaccines. The urgent need to develop active monoclonal antibodies for clinical use is obvious.

Lastly, we found that the spikes of BQ and XBB subvariants have similar binding affinities to hACE2 as the spikes of their predecessors (**Figure 5**), suggesting that the recently observed growth advantage for these novel subvariants is likely due to some other factors. Foremost may be their extreme antibody evasion properties, especially considering the extensive herd immunity built up in the population over the last three years from infections and vaccinations. BQ.1, BQ.1.1, XBB, and XBB.1 subvariants exhibit far greater antibody resistance than earlier variants, and they may fuel yet another surge of COVID-19 infections. We have collectively chased after SARS-CoV-2 variants for over two years, and each time we have fallen a step behind the evolution of the virus. This continuing challenge highlights the importance of developing vaccine and monoclonal antibody approaches that protect broadly and anticipate the antigenic trajectory of SARS-CoV-2.

## SUPPLEMENTAL INFORMATION

**Figure S1.**
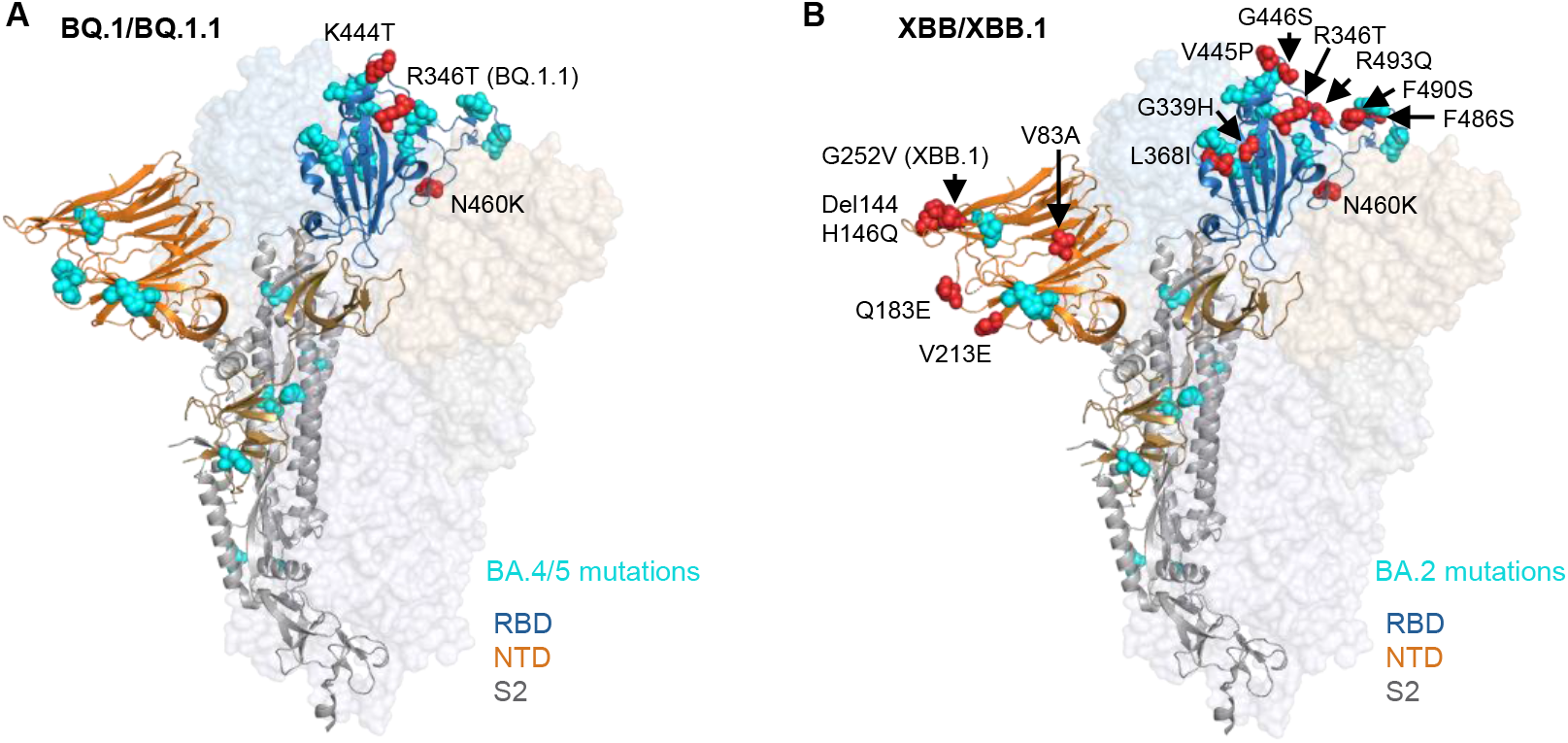
Key mutations of BQ.1 and BQ.1.1 in the context of BA.4/5 (**a**), and key mutations of XBB and XBB.1 in the context of BA.2 (**b**).

**Figure S2.**
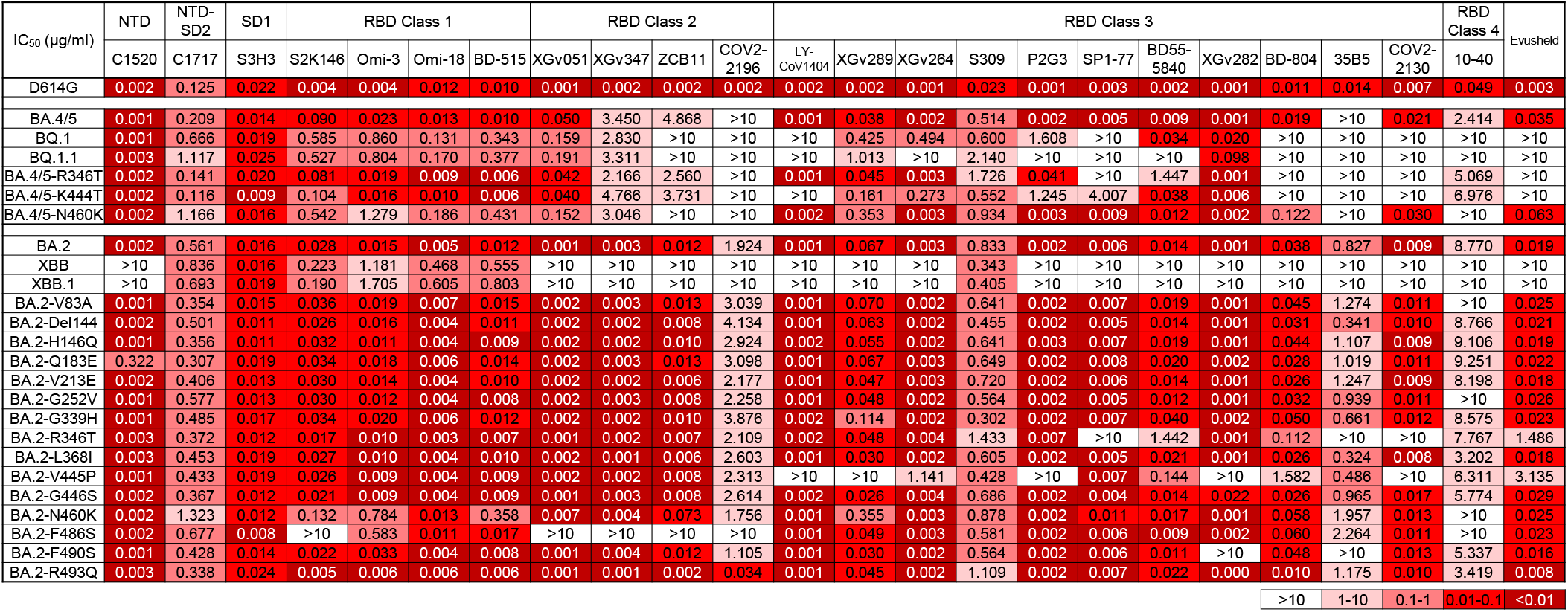
Pseudovirus neutralization IC_50_ values for mAbs against D614G, Omicron subvariants, and point mutants of BQ.1, BQ.1.1, XBB, and XBB.1 in the background of BA.4/5 or BA.2.

**Figure S3.**
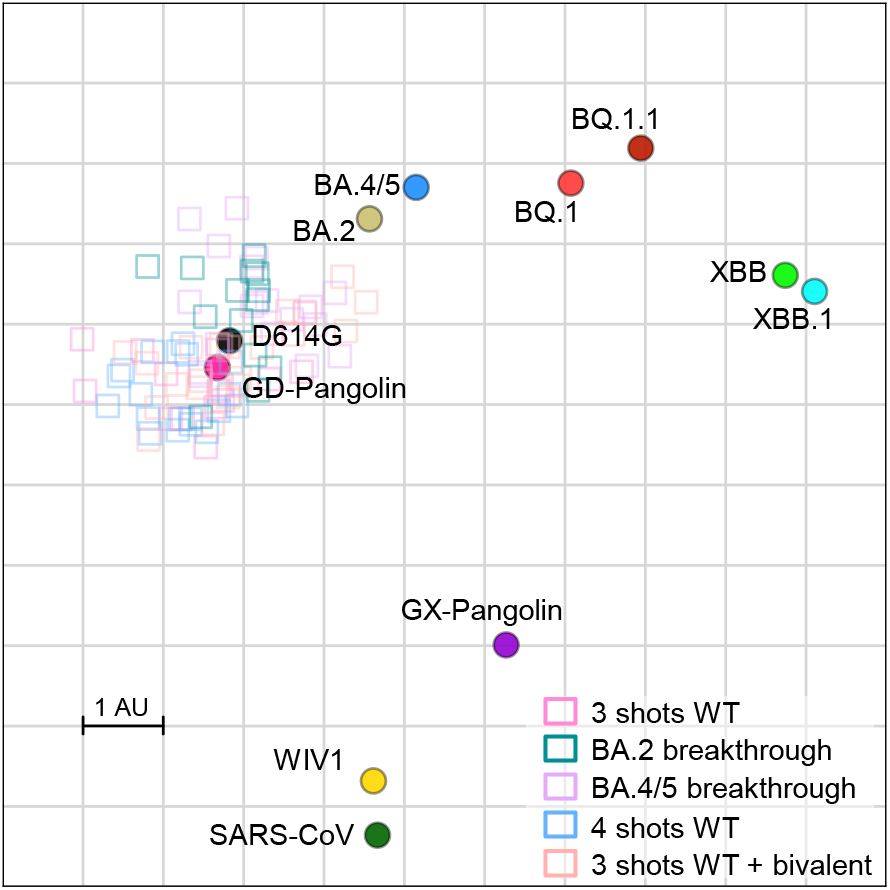
Antigenic map of BQ.1, BQ.1.1, XBB, and XBB.1 in relation to sarbecoviruses.

**Table S1.**
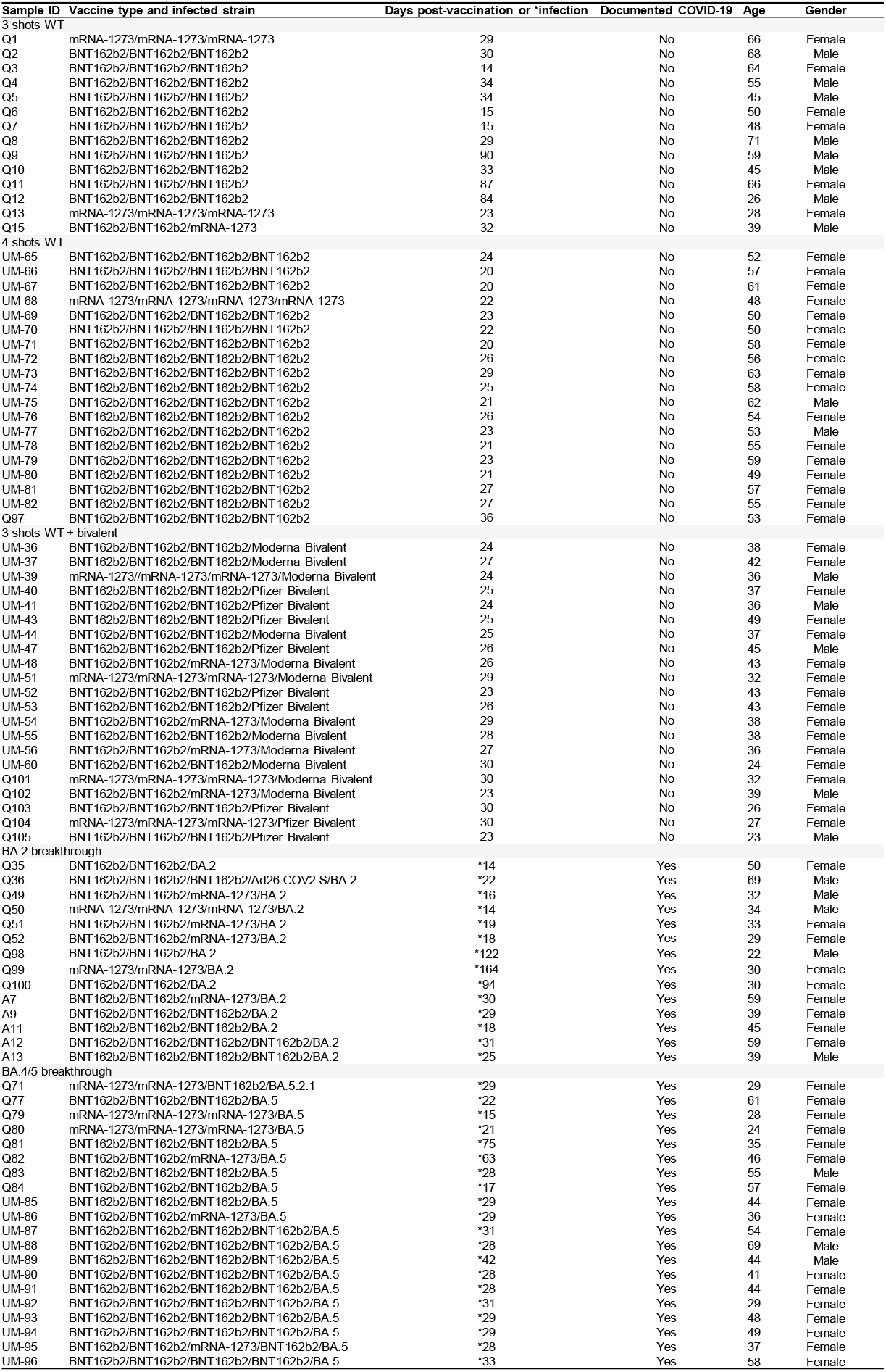
Demographics of the clinical cohorts.

## ACKNOWLEDGEMENTS

This study was supported by funding from the NIH SARS-CoV-2 Assessment of Viral Evolution (SAVE) Program and through the National Institutes of Health Collaborative Influenza Vaccine Innovation Center (75N93019C00051). We acknowledge Michael T. Yin and Magdalena E. Sobieszczyk for providing serum samples. We thank all who contributed their data to GISAID.

## AUTHOR CONTRIBUTIONS

D.D.H. and Lihong L. conceived this project. Q.W., S.I., Z.L., and Lihong L. conducted pseudovirus neutralization assays and purified SARS-CoV-2 spike proteins. Y.G. and Z.S. conducted bioinformatic analyses. Q.W., Liyuan L., Y.H., H.H.W., and Lihong L. constructed the spike expression plasmids. Q.W. managed the project. J.Y. M.W., and M.L. expressed and purified antibodies. Z.L. performed SPR assay and structural analyses. R.V., A.L., and A.G. provided clinical samples. A.B. generated antigenic map. D.D.H. and L.L. directed and supervised the project. Q.W., S.I., Z.L., Y.G., A.B., Lihong L., and D.D.H. analyzed the results and wrote the manuscript.

## METHOD DETAILS

### Human subjects

Sera analyzed in this study were categorized into several cohorts. “3 shots WT” samples were sera from individuals who had received three doses of monovalent, referred to as wild-type (WT) mRNA vaccines (either Moderna mRNA-1273 or Pfizer BNT162b2). Sera were also collected from individuals after a fourth monovalent mRNA vaccine (referred to as “4 shots WT”). Bivalent vaccine sera were collected from individuals who had received three monovalent mRNA vaccine doses followed by one dose of the Pfizer or Moderna bivalent vaccine targeting BA.4/BA.5 in addition to the ancestral D614G variant. “BA.2 breakthrough” and “BA.4/BA.5 breakthrough” sera were collected from individuals who had received monovalent mRNA vaccines followed by infection with Omicron subvariants BA.2 and BA.4 or BA.5, respectively. Samples were examined by anti-nucleoprotein (NP) ELISA to confirm status of prior SARS-CoV-2 infection. Clinical information for the different study cohorts is summarized in **Table S1**.

A subset of sera analyzed in this study was collected at Columbia University Irving Medical Center. Subjects provided written informed consent, and serum collections were performed under protocols reviewed and approved by the Institutional Review Board of Columbia University. Additional serum samples included in this study were collected at the University of Michigan through the Immunity-Associated with SARS-CoV-2 Study (IASO), which is an ongoing cohort study in Ann Arbor, Michigan that began in 2020 (Simon et al., 2022). IASO participants provided written informed consent and all serum samples were collected under the protocol reviewed and approved by the Institutional Review Board of the University of Michigan Medical School.

### Cell lines

Vero-E6 cells (CRL-1586) and HEK293T cells (CRL-3216) were purchased from the ATCC. Expi293 cells (A14527) were purchased from Thermo Fisher Scientific. Morphology of each cell line was confirmed visually before use. All cell lines tested mycoplasma negative. Vero-E6 cells are from African green monkey kidneys. HEK293T cells and Expi293 cells are of female origin.

### Monoclonal antibodies

Antibodies were generated as previously described (Liu et al., 2020). The variable regions of heavy and light chains for each antibody were synthesized (GenScript), cloned into gWiz or pCDNA3.4 vector, then transfected into Expi293 cells (Thermo Fisher Scientific) using 1 mg/mL polyethylenimine (PEI), and purified from the supernatant by affinity purification using rProtein A Sepharose (GE).

### Variant SARS-CoV-2 spike plasmid construction

Spike-expressing plasmids for D614G, BA.2, and BA.4/5 were previously generated (Wang et al., 2022d). Plasmids expressing the spike genes of BQ.1, BQ.1.1, XBB, and XBB.1, as well as the individual mutations found in the four variants in the background of BA.4/5 or BA.2 were generated by an in-house high-throughput template-guide gene synthesis approach, as previously described (Liu et al., 2022b). Briefly, 5’-phosphorylated oligo pools with designed mutations were annealed to the template of the BA.2 or BA.4/5 spike gene construct and extended by high fidelity DNA polymerase. Taq DNA ligase was used to seal nicks between extension products, which were subsequently amplified by PCR to generate variants of interest. Next generation sequencing (Baym et al., 2015) was performed on the Illumina Miseq platform (single-end mode with 50 bp R1) to verify the sequences of variants. Cutadapt v2.1 (Martin, 2011) and Bowtie2 v2.3.4 (Langmead and Salzberg, 2012) were used to analyze raw reads to get the resulting read alignments, which were then visualized in Integrative Genomics Viewer (Robinson et al., 2011).

To make the expression constructs for soluble spike trimer proteins, we subcloned the ectodomain (1-1208aa in WA1) of the spike into the paH vector and then introduced K986P and V987P substitutions as well as a “GSAS” substitution of the furin cleavage site (682-685aa in WA1) into the spike (Wrapp et al., 2020). All constructs were confirmed by Sanger sequencing.

### Protein expression and purification

To make human ACE2 protein, pcDNA3-sACE2-WT(732)-IgG1 (Chan et al., 2020) (Addgene plasmid #154104, gift of Erik Procko) plasmid was transfected into Expi293 cells using PEI at a ratio of 1:3, and the supernatants were collected after five days. hACE2 was purified from the cell supernatant by using rProtein A Sepharose (GE) followed by running through a Superdex 200 Increase 10/300 GL column. For the spike trimer proteins, paH-spike was transfected into Expi293 cells using PEI at a ratio of 1:3, and the supernatants were collected five days later. The spike proteins were purified using Excel resin (Cytiva) according to the manufacturer’s instructions. The molecular weight and purity were checked by running the proteins on SDS-PAGE.

### Surface plasmon resonance (SPR)

The CM5 chip was immobilized with anti-His antibodies using the His Capture Kit (Cytiva) to capture the spike protein through their C-terminal His-tag. Serially diluted human ACE2-Fc protein was then flowed over the chip in HBS-EP+ buffer (Cytiva). Binding affinities were measured with the Biacore T200 system at 25_□ in the single-cycle mode. Data was analyzed by the Evaluation Software using the 1:1 binding model.

### Pseudovirus production

SARS-CoV-2 pseudoviruses were generated as previously described (Liu et al., 2020). In brief, HEK293T cells were transfected with a spike-expressing construct using 1 mg/mL PEI and then infected with VSV-G pseudotyped ΔG-luciferase (G*ΔG-luciferase, Kerafast) one day post-transfection. 2 hours after infection, cells were washed three times with PBS, changed to fresh medium, and then cultured for one more day before the cell supernatants were harvested. Pseudoviruses in the cell supernatants were clarified by centrifugation, aliquoted, and stored at -80°C.

### Pseudovirus neutralization assay

Pseudoviruses were titrated on Vero-E6 cells before conducting the neutralization assays to normalize the viral input between assays. Heat-inactivated sera were serially diluted starting from 1:100 with a dilution factor of four and antibodies were 5-fold serially diluted starting from 10 µg/mL in 96 well plates in triplicate. Then, 50 µL of diluted pseudovirus was added and incubated with 50 µL serial dilutions of serum or antibody for 1 hour at 37°C. During the co-culture, Vero-E6 cells were trypsinized, resuspended with fresh medium, and then added into virus-sample mixture at a density of 4□×□10^4^ cells/well. The plates were incubated at 37°C for ~12 hours before luciferase activity was quantified using the Luciferase Assay System (Promega) using SoftMax Pro v.7.0.2 (Molecular Devices). Neutralization ID_50_ values for sera and IC_50_ values for antibodies were calculated by fitting a nonlinear five-parameter dose-response curve to the data in GraphPad Prism v.9.2.

### Antibody footprint and mutagenesis analysis

All the structures were downloaded from the Protein Data Bank (7XIV (BA.2 spike), 7WK9 (S3H3), 7UAR (C1717), 7UAP (C1520), 7TAS (S2K146), 7XCO (S309), 7WRZ (BD55-5840), 7ZF3 (Omi-3), 7ZFB (Omi-18), 7E88 (BD-515), 7WED (XGv347), 7XH8 (ZCB11), 7SD5 (10-40), 7WM0 (35B5), 7WLC (XGv282), 7WE9 (XGv289), 7UPY (SP1-77), 7QTK (P2G3), 7MMO (LY-CoV1404), 7EYA (BD-804)) for analysis. The interface residues were obtained by running the InterfaceResidues script from PyMOLWiki in PyMOL, and the edge of these residues was defined as the footprint of the antibodies. Site-directed mutagenesis was also conducted in PyMOL. All the structural analysis figures were generated in PyMOL v.2.3.2 (Schrödinger, LLC).

### Antigenic cartography

We constructed an antigenic map based on the serum neutralization data by utilizing the antigenic cartography technique as previously described (Wilks et al., 2022). The antigenic map was generated using the Racmacs package (https://acorg.github.io/Racmacs/, version 1.1.35) in R with 1000 optimization steps, a dilution step size of zero, and the minimum column basis parameter set to “none”. All distances between virus and serum positions on the antigenic map were optimized such that distances correspond to the fold decrease in neutralizing ID_50_ titer, relative to the maximum titer for each serum. Each unit of distance in any direction in the antigenic map corresponds to a two-fold change in the ID_50_ titer.

### Quantification and statistical analysis

IC_50_ and ID_50_ values were determined by fitting the data to five-parameter dose-response curves in GraphPad Prism v.9.2. Comparisons were made by two-tailed Wilcoxon matched-pairs signed-rank tests. \*\*\**p* < 0.001; \*\*\*\**p* < 0.0001.

